# Mechanical Behavior of Axonal Microtubules; the Effect of Fluid on the Rupture of Axonal Microtubules

**DOI:** 10.1101/378455

**Authors:** Farid Manuchehrfar, Amir Shamloo

## Abstract

Axonal microtubules are dynamically instable bundles in the interior part of the axon. The dynamics of these bundles are of vital importance in the behavior of axon such as their degeneration. Each axon typically contains 10~100 microtubule bundles with average length of 4μm. These bundles are coated with cytoplasm and are cross linked with random number of tau proteins. In some circumstances such as acceleration or deceleration of head in space or during the strike, they are placed in tension which may cause rupture of these bundles or disconnection of tau protein cross links. Mechanical behavior and rupture modality of microtubule bundles are becoming more and more important recently. In our model, viscoelastic microtubule bundles constituted from several discrete masses connected to the neighboring mass with a standard linear solid (SLS), a spring damper model. In addition we take into account the effect of cytoplasm by Dissipative Particle Dynamic (DPD) to investigate the rupture nature and mechanical behavior of these bundles and the effect of cytoplasm on their mechanical behavior. We obtain these results for various amounts of suddenly applied end forces to the group of axonal microtubule bundles.

## 1. INTRODUCTION

Axonal microtubules are dynamically instable and microtubules cycle through the length of axons. Dynamic instability of microtubules causes the microtubules to occupy greater volume in the interior of axons (Buxton et al. 2010). Microtubules’ dynamics are impressive in the function of axons and their degeneration (Buxton et al. 2010). Microtubular geometry is formed by microtubule-associated (Tau) proteins and tubulin polymerization (Shamloo et al. 2016).

Microtubular function can be affected by acetylation, glutamylation, glycation, phosphorylation and tyrosination (Perez et al. 1999, Shamloo and Mehrafrooz 2018) and dysfunctions of axonal microtubules can result in Alzheimer’s disease (AD) neurons (Majid T. et al. 2015). In AD neuron, microtubules reduce in number and length, significantly (Buxton et al. 2010).

Typically each axon contains ~10-100 microtubule/cross sectional view (Fadic et al 1985) and the average length of each microtubule bundle is ~4μm in the axon (Yu et al. 1994) and they have polar orientations. Axonal microtubules are cross linked to their neighboring microtubule bundles with tau proteins and these tau proteins are considered as one of the important elements for axonal strength in response to the mechanical force (Prechsel et al. 1992).

In some circumstances, such as in acceleration or deceleration of head in the space, microtubule bundles are places under tension force (Tang-Schomer et al. 2010). The tension applied to the nervous system cause Traumatic Axonal Injuries. Axonal injury is one of the most common nervous system injuries (Smith et al. 2003, Smith et al. 2010, Sanjith 2011, Kirkcaldie et al. 2016) and patients who are suffering from axonal injuries are commonly experience numerous functional deficits (Adam et al. 1989, Sanjith 2011, Blumberg et al. 1989, Williams et al. 1990).

Microtubule bundles can resist mechanical stress (Peter and Mofrad 2012). Since the injuries of microtubules are shown to be one of the main consequences of the traumatic brain injury these bundles have been studied widely from mechanical point of view (Margulies et al. 1990, Shamloo et al. 2015A). Most studies also show that the rupture of microtubules causes severe damages in axonal brain injuries (Tang-Schomer et al. 2010, Mehrafrooz and Shamloo 2018).

There are numerous models which study the mechanical injuries of microtubules computationally and experimentally (Kilinc et al. 2008 and 2009, Meaney et al. 1994, Dixton et all 1988 and 1991, Feeney et al. 1991, Mishra et al. 2005). Mathematical and computational modeling has been proved to be the easiest way to study the mechanics of nervous system (Erickson and O’Brein 1992, Flyvbjerg et al, Shamloo et al. 2015 B and C). In axonal computational modeling, microtubule bundles are assumed to be composed of discerete masses connected with mechanical elements (Buxton et al. 2010, Wu et al. 2018).

Microtubule bundles have been recently modeled both as discrete masses and continuum elements (Shahinnejad et al. 2013, Teixeira et al. 2014, Ansari-Benam et al. 2015, Wu et al. 2018, Adnan et al. 2018). Shahinnejad et al. 2013 used finite volume to model axonal microtubules under tension and they also assumed microtubules as discrete masses. In 2012, Peter and Mofrad suggested a model for axonal microtubules. They endeavored to represent an intact model for axonal microtubules. In their model microtubules were assumed to be lots of discrete masses connected to their neighboring microtubule bundle with number of tau proteins (a system of spring and damper), microtubule bundles are positioned in a hexagonal array and each microtubule bundle was considered to have one point of discontinuity. Microtubule bundles were assumed to have the average length of 8μm in their model (Soheilypour et al. 2015). Another study in 2014 modeled different microtubules and their connected tau proteins under the specific amounts of strain and checked the effect of microtubule length on their mechanical behavior (Ahmadzadeh et al. 2014).

Microtubules are coated with fluids and cytoplasm which affects mechanical behavior of axonal microtubules. Interaction of fluid particles and microtubule bundles changes the mechanical response of microtubule comparing to their behavior in the absence of cytoplasm. In our model, the effect of fluid around the microtubules on the behavior of microtubules under the action of varying stresses is studied in a three dimensional dimensional (2D) platform. We determined the steady state behavior of axon for different force rates and different force magnitudes under the effect of cytoplasm and compared this behavior with one which is not under the effect of cytoplasm. In Fig. 1(a), a neuronal cell is shown with its connected axon, dendrites and microtubules. As it is shown in this figure, the axonal microtubules are positioned within the interior part of axon (Fig. 1(b)).

**Fig. 1.**
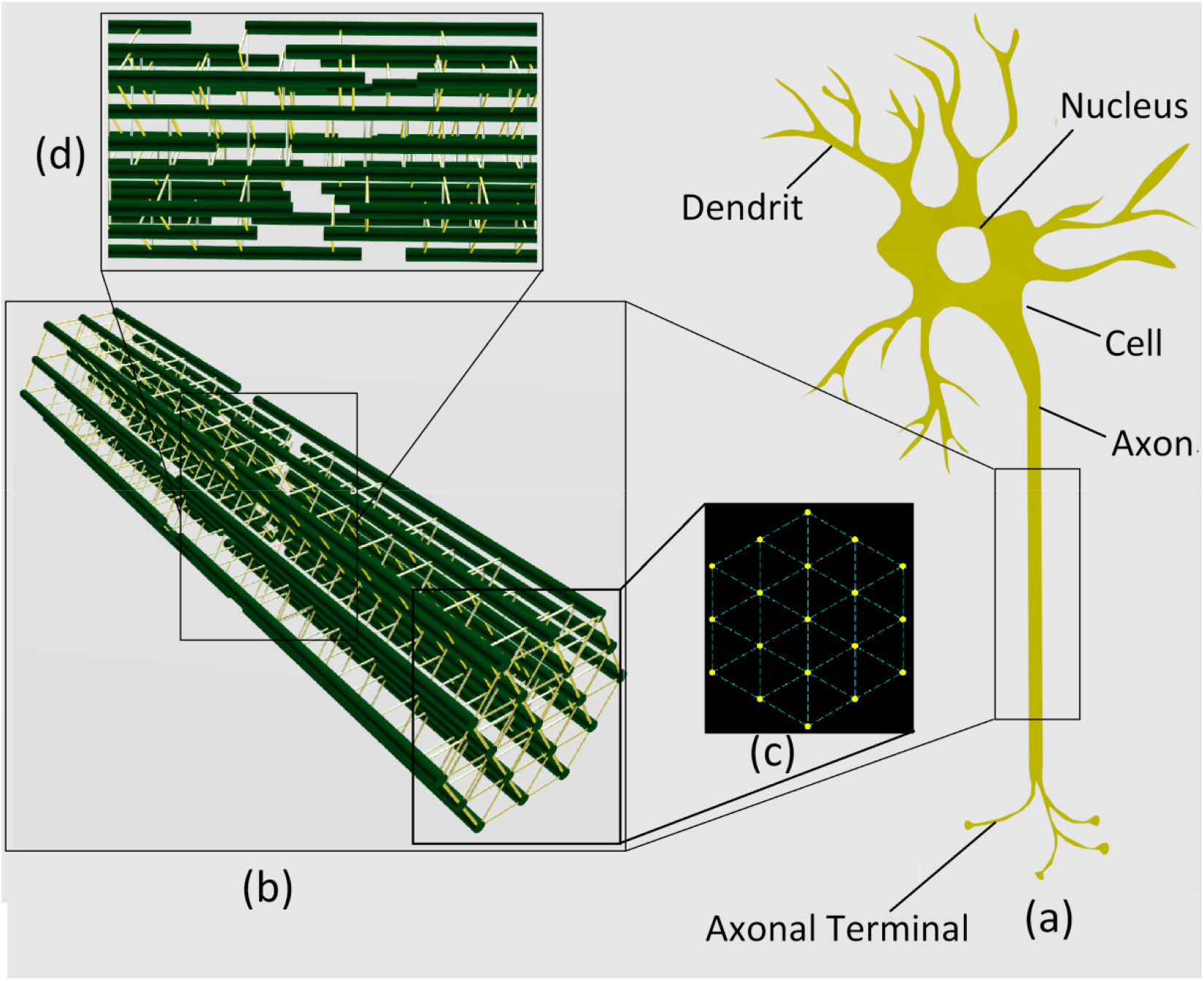
Schematic of a neural cell in part (a) with its axon, dendrites and axonal terminals; in part (b) we see our 3 dimensional platform built with 19 microtubule bundles connected to each other in hexagonal cross section (part (c)); microtubule bundles are cross linked to each other with tau proteins (Yellow cross links in part (d)) and each microtubule has a point of discontinuity (part(d)).

## 2. MODELING

### a. Geometry of Microtubule Bundles

In this study, axonal microtubules are assumed to be hexagonal array of 19 bundles in a three dimensional platform (Fig. 1(b)&(c)) (Peter and Mofrad 2012, Shamloo et al. 2015A). The length of each microtubule bundle is 8μm. We model each microtubule bundle as ensemble of 800 discrete masses connected to their neighboring mass by a Standard Linear Solid (SLS) Unit. Fig.2 is a schematic to show how discrete masses connect to their neighbor with SLS units. One point of discontinuity on each bundle is considered by eliminating one of the SLS units within middle 80% of the microtubule bundle (Fig. 1(d)). Microtubule bundles are cross linked to their neighboring microtubule through Tau proteins. In addition, between each two neighboring microtubules a random number of cross linking tau proteins exist which their locations are also on a random basis (Shamloo et al. 2015A). In Fig. 1(c) a cross sectional view of Three-Dimensional microtubule bundle is represented and we can observe a zoomed view of bundles cross linked with tau proteins in Fig. 1(d). The space between each two microtubule bundle is 0.045 μm which is 4.5 times the distance between each two neighboring mass on an individual bundle which is filled with cytoplasm.

**Fig. 2.**
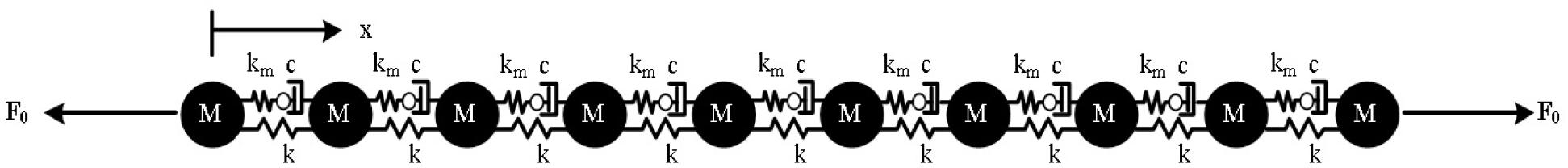
10 discrete masses connected to their neighbors with a SLS unit; to show how we connect discrete masses in an individual microtubule bundle

### b. Fluid Modeling

Fluid around the microtubule bundles is modeled using Dissipative Particle Dynamic (DPD) method (Liu et al. 2014). Fluid is assumed to be a large number of DPD particles which filled the whole surrounding of axonal microtubules. The allocation of DPD particles is face-centered cubic lattice which is dense enough to model fluid particles. To model the random position of these particles, we changed the initial position of each particle randomly by a very small ratio (maximum of 0.1 of distance between each 2 particles). In addition, Maxwell-Boltzmann distribution has been used in this study to specify the initial velocity of each DPD particle (Walstad 2013). Fig. 3 illustrates 3 dimensional axonal microtubules surrounded with fluid particles.

**Fig. 3.**
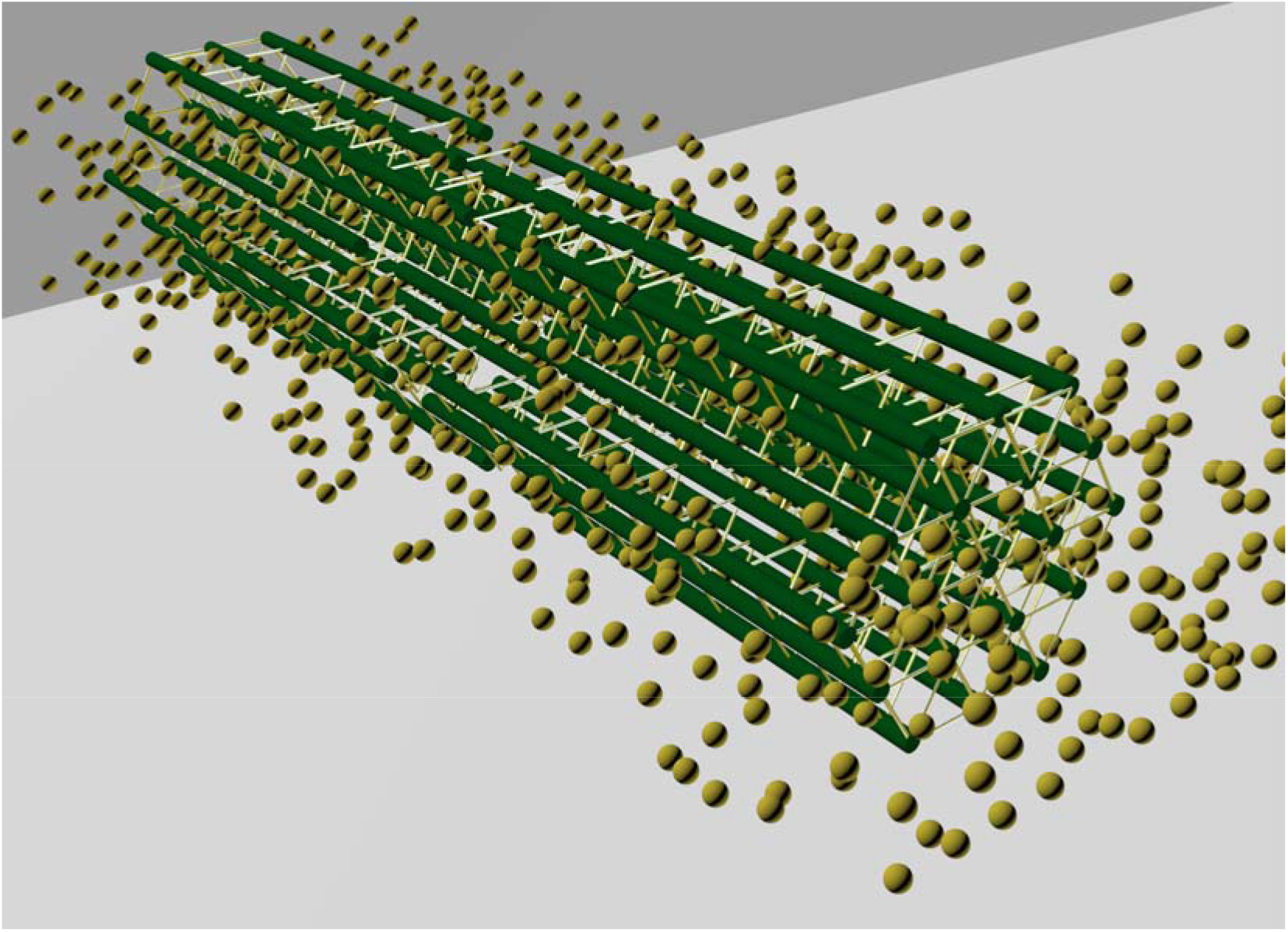
Axonal microtubules cross linked with tau proteins and surrounded with fluid particles (yellow particles)

### c. Forces

Dissipative Particle Dynamic (DPD) follows the basic principles of Navier-Stokes equations with certain alteration (Liu et al. 2006). The computational cost of this method is much less than other similar methods such as Molecular Dynamic (MD) (Keaveny et al. 2005). DPD forces are considered all possible inter-particle forces between fluid-fluid particles and fluid-Microtubule bundle particles There are three DPD forces exerted on particle ‘i’ by surrounding particle ‘j’ (Eq. 1): Conservative Repulsive Force (F^C^_ij_), Dissipative Force (F^D^_ij_), and Random Force (F^R^_ij_)

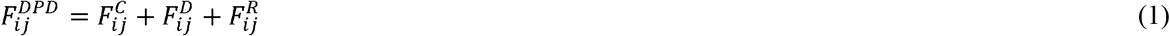

Conservative Repulsive Force is given by Eq.2. In this equation ‘a_ij_’ is the maximum repulsion force between two particles ‘i’ and ‘j’, ‘r_ij_’ is the distance between particle ‘i’ and particle ‘j’, 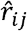 is the unit vector that determines the direction from particle ‘j’ to ‘i’. The Conservative repulsive force depends on the cut-off radius (r_c_) which is the maximum range of F^C^.

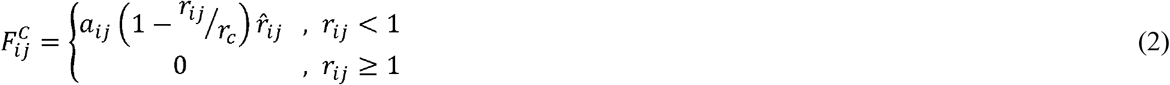

In addition, the dissipative and random forces are given by the equations 3 and 4, respectively, where ‘γ’ and ‘σ’ are characteristic strength of these forces and their relation are given by equation 5.

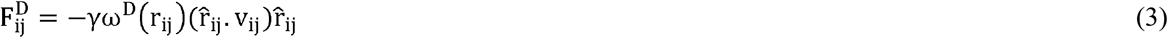

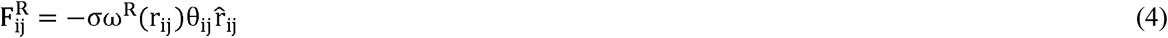

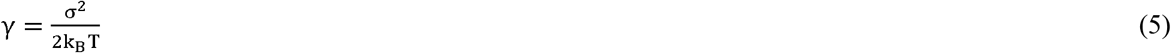

In these equations ‘V_ij_’ is the relative velocity of particle ‘i’ related to particle ‘j’, θ_ij_ is a random value with zero mean and unit variance, ω^R^ and ω^D^ = (ω^R^)^2^ are weighting functions of dissipative and random forces, respectively and can be defined by eq. 6, ‘k_b_’ is the Boltzmann’s constant and ‘T’ is the Temperature of the system.

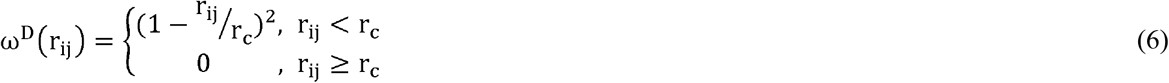

It can be considered that each particle can be affected by 7 effective forces: all possible external forces (F^ext^_ij_); force between 2 neighboring mass in a same microtubule bundle (F^SLS^_ij_); force that is acted upon tau proteins by a neighboring mass from the neighboring microtubule bundle (F^Tau^_ij_); the force which is due to the effect of bending moment of a sole microtubule bundle (F^b^_ij_); force between a DPD particle and a discrete mass of microtubule bundle which is known as particle-microtubule bundle force (F^P-M^_ij_); and force between two DPD particles known as particle-particle force (F^P-P^_ij_) and as it is noted earlier ‘F^P-M^_ij_’ and ‘F^P-P^_ij_’ are themselves divided into 3 separately part: Conservative repulsive force, dissipative force, and random force. SLS forces also contain 2 parts: elastic force which is because of the springs in a SLS unit, ‘F^el^_ij_’, and dissipative force of damper in the SLS system, ‘F^P-P^_ij_’ 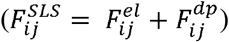.

We have written a MATLAB code to solve our system. In our code each particle and microtubular discrete mass has an index which make particle known during various iterations. In each time step and for each particle we measure the total force from other particles as in equation 7. When the force of a particle is measured, we are able to determine the acceleration of the particle; as a result the velocity and location of the particle in the next time step can be define using numerical integrating and derivation methods. In equation 7, ‘a’, ‘b’, ‘c’ and ‘d’ are indicators of whether particle ‘i’ and particle ‘j’ are discrete masses in the same microtubule; discrete masses in 2 various neighboring microtubule; a fluid particle and a discrete mass of a microtubule bundle; or 2 fluid particle. These indicators are logical numbers and assume only binary values of unity or zero. To illustrate it more, if the particle ‘i’ is a discrete mass of a microtubule bundle and particle ‘j’ is a DPD particle, there would be exist only 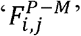 and ‘d’ would be zero numbers. We let the system goes until it reach its steady state. In various time steps, if the strain of each two neighboring mass of microtubule bundles which are connected together with SLS unit become more than 50%, the SLS unit is fractured and when the fractures increase, the system breaks down and the rupture occurs.

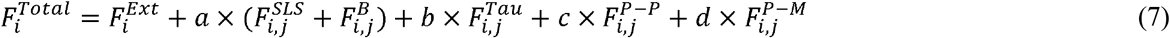

## 3. RESULT

We have modeled axonal microtubule bundles computationally and we are interested in finding the mechanical behavior of them under the effect of suddenly applied end forces in the presence of the fluid around them and compare our result with their behavior in the absence of fluid to find the effect of fluid on their mechanical behavior.

In the beginning, we make all of our equations non-dimensional. For finding the appropriate time step we used 10 discrete masses connected to each other with SLS units (Fig. 2). Using the system we found the appropriate time step will be equal to ~ 2.245×10^(−2)^ ps.

Table 1 shows the material parameters used in this study (Peter and Mofrad 2012). If the microtubule bundles shown in Fig. 1(b) are assumed to be attached to a constant wall in one side (Cell body side) and act upon on other side (Axonal terminal side) by an external force, microtubule bundles will show time-varying deformation. When bundles start deforming, the overall strain of the system in each time step can be defined by Eq. (8). The first and the second summation in this equation are the over the rightmost and leftmost points. ‘n’ and ‘L*’ are the number of microtubules and the initial non-dimensional length of the system.

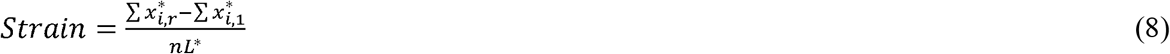

**Table1.** Material parameters used in our computational platform

| Parameter | Value |
| --- | --- |
| Microtubule Young's modulus | 1.5 GPa |
| Microtubule Persistence Length | 420 μm |
| Microtubule Flexural Rigidity | $1.8 \times 10^{-24}$ Nm<sup>2</sup> |
| Tau Protein Young's Modulus | 5 Mpa |
| Microtubule Element Length | 10 nm |
| Tau Protein Element Length | 45 nm |
| Microtubule Axial Spring Constant | 47.1 N/m |
| Microtubule Bending Spring Constant | $1.8 \times 10^{-16}$ Nm |
| Tau cross-link axial spring constant | $3.925 \times 10^{-2}$ N/m |
| Microtubule bead mass | $1.48375 \times 10^{-21}$ Kg |
| Tau cross-link bead mass | $2 \times 10^{-22}$ Kg |
| Microtubule Damping Ratio | 2.1191 |
| Tau Protein Damping Ratio | 5.9555 |

According to Janmey et al (1991), we assumed that when the strain between two neighboring point mass of microtubule bundles become more than 50%, the SLS unit between them breaks down.

During the real traumatic brain injury, the mechanical force on the neurons can be considered to grow rapidly in the time interval of [0, τ] (‘τ’ is in the nanoseconds scale.) from zero to its final value (F) and then become a constant value in the further time steps (Fig.4). We investigated the response of our system of microtubules for 5 values of non-dimensional applied force, ‘F’, in the interval of [0.1, 0.275] (between 0.0471 and 0.122 mN).

**Fig. 4.**
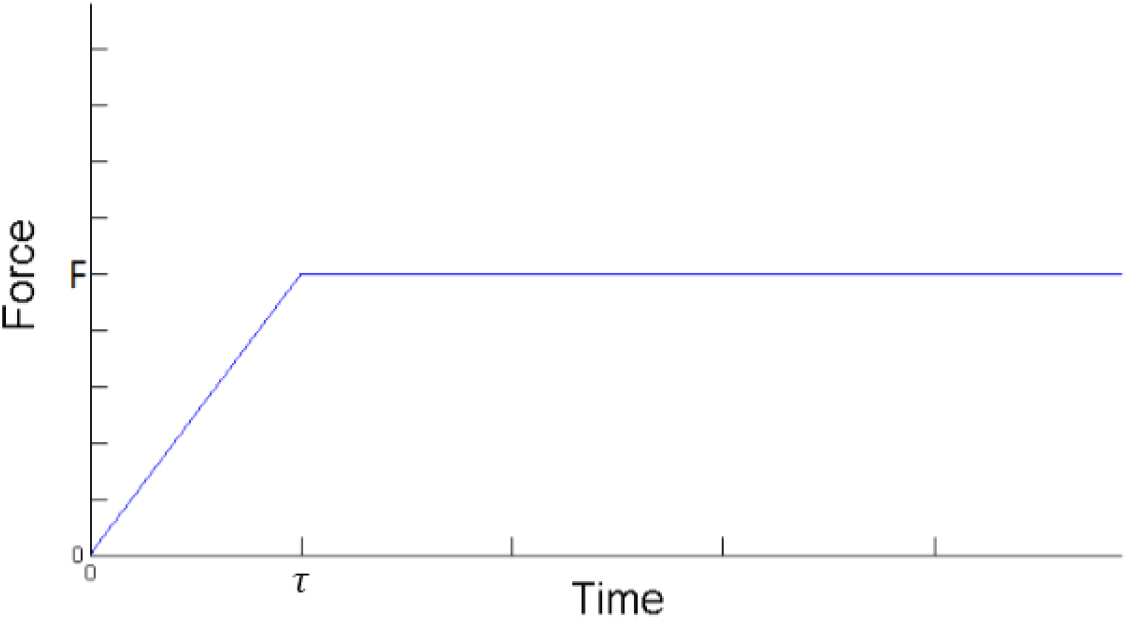
Applied force to the end of microtubule bundles; force grows linearly in a growing time *τ* and after that becomes constant.

Fluid around the microtubules is expected to dissipate the energy of system like dampers, as a result it is expected that the microtubule bundles are safer in the present of the fluid around them due to the energy dissipation of fluid. Thus, in the presence of the fluid the steady strain of the system of microtubules are decreased. Fig. 5 represents the Strain-Time diagram of microtubules in the presence of fluid around them with the system which is not in the contact of fluid for various values of ‘F’. (‘τ’ is equal to 15000dt). The value of non-dimensional ‘F’ in Fig. 5(a) is 0.1 (4.71 μN). We also studied the effect of fluid for two other acting forces (0.15 and 0.2), which is shown in Fig. 5(b) & (c). In Fig. 5, the dashed lines represent the dynamic response of axonal microtubule bundles when we do not consider the effect of discontinuities. Solid red lines shows their behavior when we do include the effect of discontinuities but the effect of fluid is not considered, and finally the solid blue lines represent the strain-time diagram of axonal microtubules when we consider the fluid around them and model them using DPD.

**Fig. 5.**
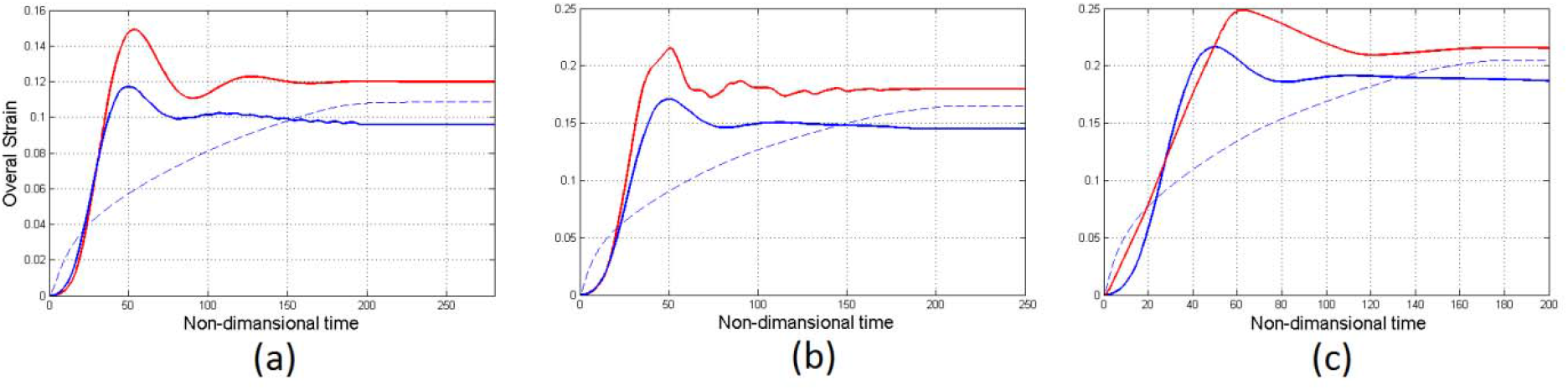
Dynamic response of axonal microtubules for three different forces (a) F=4.71 μN; (b) F=7.065 μN; (c) F=9.42 μN, in 3 different conditions: Blue dashed line in when we do not take into account discontinuity of microtubules and we assume there is no fluid around microtubules; Red line is when we consider discontinuity in each microtubule bundle and we assume there is no fluid; and blue line is when we assume both fluid and discontinuity of microtubules

It can be inferred from the Fig. 5 that when the effect of discontinuities is modeled, the steady strain of the system is increased. We could point out that the discontinuities cause growth of the system’s strain. When the surrounding fluid is modeled, the steady state strain of the system is decreased significantly and this shows that fluid decrease the stored energy of the system. According to this figure and our modeling we conclude that the fluid around the microtubules cause decreasing of final strain by 12% and this causes postponing the rupture of axonal system.

Fig. 6 indicates dynamic response of axonal microtubules for 5 varying forces, 0.1, 0.15, 0.2, 0.25, 0.275 for *τ* = 15000*dt* (84.2ns). As it can be observed from this figure, the strain is increased by increasing the applied force. Assuming the initial geometry of microtubule bundles to be the one shown in the Fig 3, which is coated with fluid particles, when the force is applied to the system, the deformation starts until the system reaches its steady state. As we increase the applied force, the steady state strain is increased and when the final non-dimensional applied force reaches its critical value of 0.275 (~ 13 μN) the rupture occurs and the connection between neuron cell and axonal terminals breaks down.

**Fig. 6.**
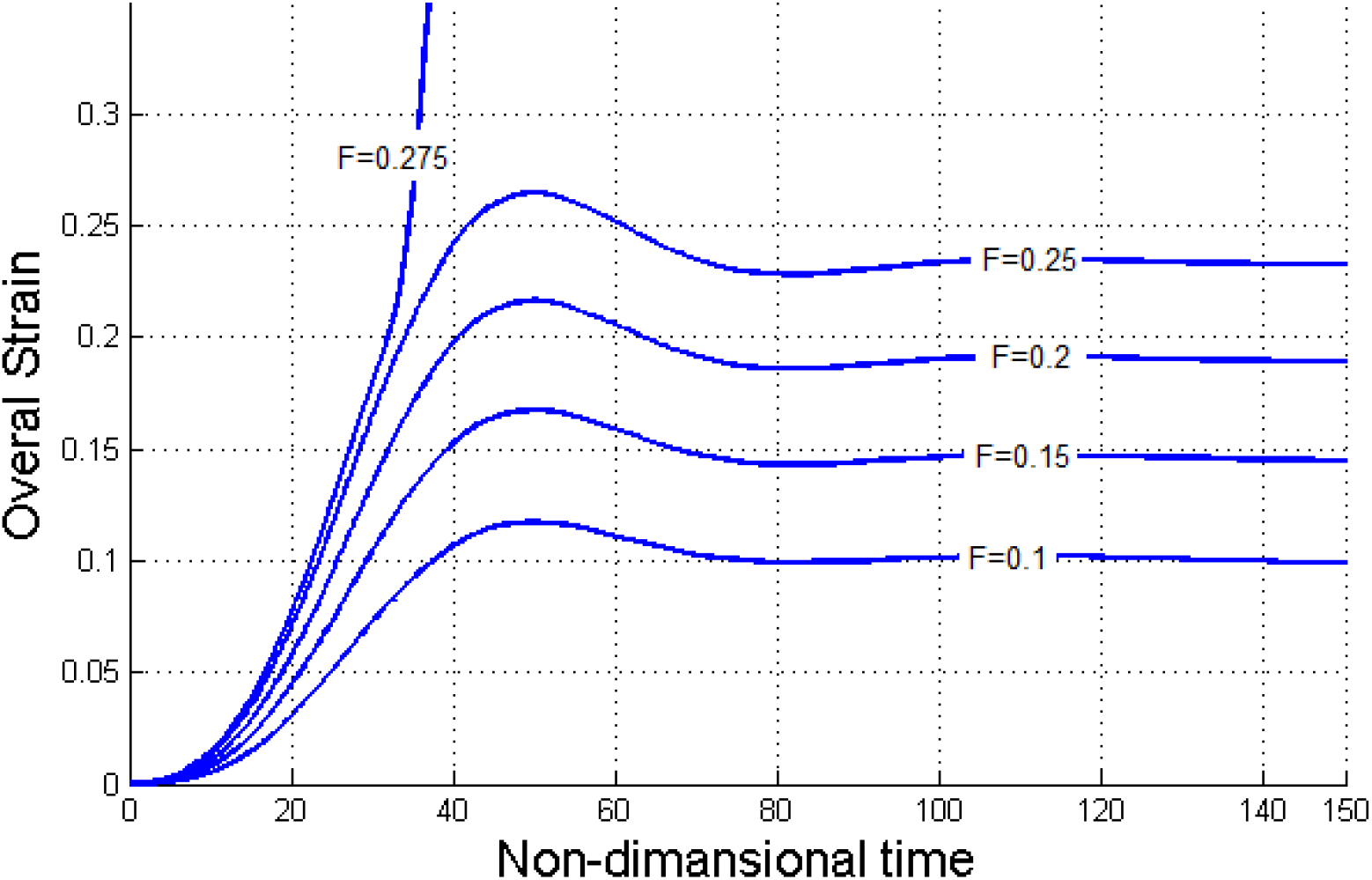
dynamic response of axonal microtubules for 5 varying forces, 0.1, 0.15, 0.2, 0.25, 0.275 for *τ* = 15000dt (84.2ns)

We studied the nature of rupture in this study, including how rupture starts, expands and failure occurs. Fig. 7 illustrates the nature of rupture when the non-dimensional acting force becomes 0.275 (~13μN). The bottom part in this figure shows the initial position of microtubule bundles and the location of discontinuities. When the deformation starts and the strain increases gradually, as the force is high enough to breaks the connections down, rupture begins at the non-dimensional time of about ~ 30 (0.173ns) and outspreads rapidly and finally at the non-dimensional time of about 40 the failure happens and no signal can be transferred from neuronal cell body to axonal terminals (top figure in Fig 7).

**Fig. 7.**
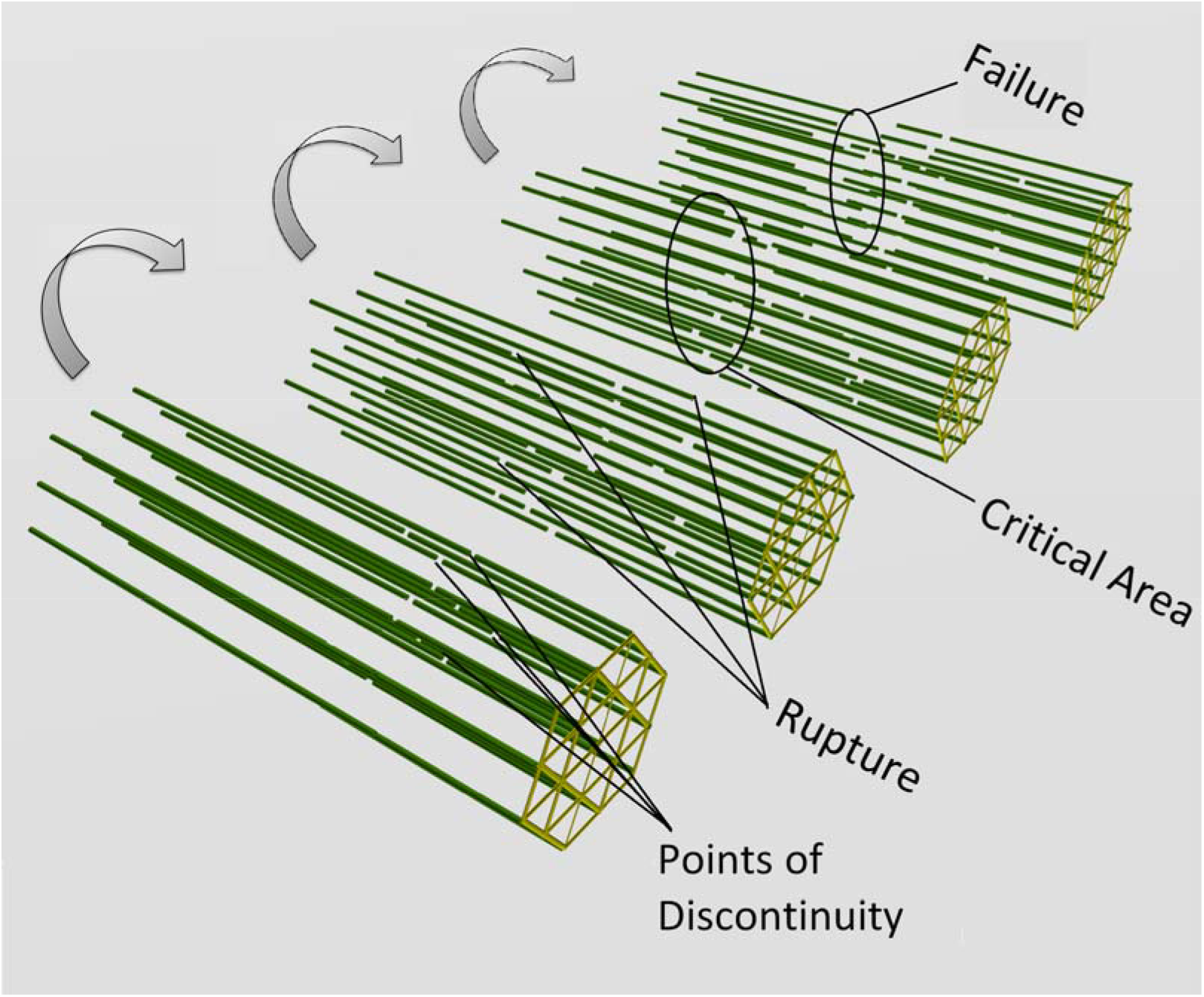
Failure of axonal microtubules when the non-dimensional acting force becomes 0.275 (~13μN)

We studied the effect of different parameters on the critical area of axonal microtubule bundles, where bundles are more likely to fail. Based on our observation over various forces and different force-growing times (**τ** in Fig. 4), critical area is always nearer to the axonal terminal compared to its distance from cell body (Fig.8). When we increase **τ** (force grows slowly) while force is over-critical, the critical area moves to the middle of the axonal microtubules and when τ is decreased, critical area becomes nearer to the axonal terminals.

**Fig. 8.**
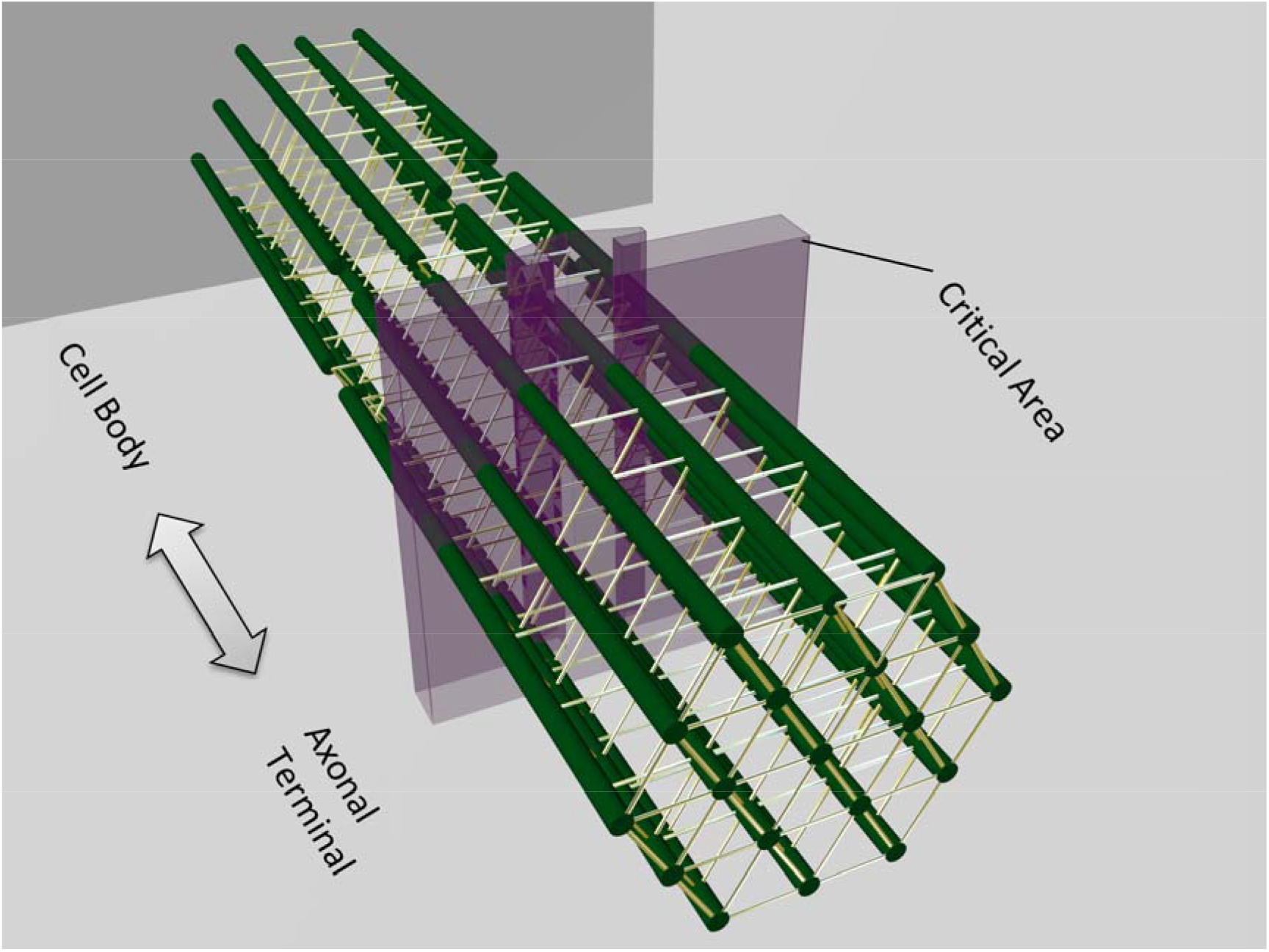
Critical area of axonal microtubules, when force-growing time (τ in Fig. 4) decreases, critical area shows tendency to become nearer to axonal terminals

## 4. CONCLUSION

Knowing the exact behavior of axonal microtubules is very helpful in investigating the damage on central nervous system. We modeled axonal microtubules computationally using Spring-Damper methods. We assumed microtubule bundles as a series of discrete masses connected to their neighbors with SLS units while each bundle has a point of discontinuity. Surrounded cytoplasm of bundles is modeled with Dissipative Particle Dynamic (DPD) method and interaction of microtubule bundles with fluid particles are considered to simulate the mechanical response of axonal microtubules to a specific external forces. These forces were acted to the end of each microtubule bundle.

Now, we are able to predict the nature of axonal rupture and failure of axonal microtubules and we identified the critical damage areas of microtubules. We modeled microtubules which are relatively close to the real behavior of axonal microtubules, comparing with other models. We demonstrate that the failure of axonal microtubules occur when the applied force become about 13μN.

The effect of cytoplasm on axonal microtubules is studied in our model. It is indicated that fluid around the microtubules dissipate the energy of system and cause the strain to decrease by 12% and as a result decrease the feasibility of mechanical failure of axonal microtubules which are responsible to transfer the information from cell body to axonal terminals.

Computational modeling of axonal microtubules is very important in knowing the behavior of microtubules due to the fact that it is not only time-efficient, but also easy and cost-effecting. Mechanical damage of neurons is one of the most common central nervous system injuries and microtubular failure is one of the main reasons of traumatic brain injury. Using the exact information of computational modeling of axonal microtubules assists us to develop new methods to prevent the damage on nervous system and culture microtubule bundles and replace the damaged part of real microtubules with cultured ones. In this model we improved other models by taking into account the effect of cytoplasm on the behavior and mechanical response of axonal microtubules.

## 5. Acknowledgement

The authors would like to thank Sharif University of Technology research center and the National Elite Institute for supporting this study.

## References

Adams, JH., Doyle, D., Ford, I., Gennarelli, TA., Graham, DI., McLellan, DR., 1989. Diffuse axonal injury in head injury: definition, diagnosis and grading. Histopathology 15, 49–59.

Adnan, A., Qidwai, S., Bagchi, A., 2018. On the atomistic-based continuum viscoelastic constitutive relations for axonal microtubules. Journal of the Mechanical Behavior of Biomedical Materials 86, 375–389. doi:10.1016/j.jmbbm.2018.06.031

Anssari-Benam, A., Bucchi, A., Bader, D.L., 2015. Unified viscoelasticity: Applying discrete element models to soft tissues with two characteristic times. Journal of Biomechanics 48, 3128–3134.

Blumbergs, P. C., Jones, N. R., North, J. B., 1989. Diffuse axonal injury in head trauma. Journal of Neurology, Neurosurgery, and Psychiatry 52 838–841.

Buxton, G.A., Siedlak, S.L., Perry, G., Smith, M.A., 2010. Mathematical modeling of microtubule dynamics: Insights into physiology and disease. Progress in Neurobiology 92, 478–483.

Dixon, CE., Lighthall, J.W., Anderson, T.E., 1988. Physiologic, histopathologic, and cineradiographic characterization of a new fluid percussion model of experimental brain injury in the rat. Journal of Neurotrauma 5 91–104.

Drechsel, D.N., Hyman, A.A., Cobb, M.H., Kirschner, M.W., 1992. Modulation of the dynamic instability of tubulin assembly by the microtubule-associated protein tau. Mol.Biol.Cell3, 1141–1154.

Erickson, H.P., O’Brien, E.T., 1992. Microtubule dynamic instability and GTP hydrolysis. Annual Review of Biophysical and Biomolecular Structure 21, 145–166.

Fadic, R., Vergara, J., Alvarez, J., 1985. Microtubules and caliber of central and peripheral processes of sensory axons. Journal of Comparative Neurology 236, 258–264.

Feeney, D., Boyeson, M., Linn, R., Murray, H., Dail, W., 1981. Responses to cortical injury: Methodology and local effects of contusion in the rat. Journal of Brain Research 211, 67–77.

Flyvbjerg, H., Holy, T.E., Leibler, S., 1994. Stochastic dynamics of microtubules: a model for caps and catastrophes. Physical Review Letter 73, 2372–2375.

Hossein Ahmadzadeh, Douglas H. Smith, Vivek B. Shenoy, 2014. Viscoelasticity of tau proteins leads to strain rate dependent breaking of microtubules during axonal stretch injury: predictions from a mathematical model. Biophysical Journal 106, 1123–1133

Janmey, P.A., 1991. Viscoelastic properties of vimentin compared with other filamentous biopolymer networks. The Journal of Cell Biology 113, 155–160.

Keaveny, E.E., Pivkin, I.V., Maxey, M., Karniadakis, G.E., 2005. A comparative study between dissipative particle dynamics and molecular dynamics for simple-and complex-geometry flows. The Journal of Chemical Physics 123, 104107.

Kilinc, D., Gallob, G., Barbee, K. A., 2009. Interactive image analysis programs for quantifying injury-induced axonal beading and microtubule disruption, computer methods and programs in biomedicine 95, 62–71.

Kilinc, D., Gallo, G., Barbee, K.A., 2008. Mechanically induced membrane portion causes axonal beading and localized cytoskeletal damage. Experimental Neurology 212, 422–430.

Kirkcaldie, M.T., Collins, J.M., 2016. The axon as a physical structure in health and acute trauma. Journal of Chemical Neuroanatomy 76, 9–18.

Liu, M., Meakin, P., Huang, H., 2006. Dissipative particle dynamics with attractive and repulsive particle-particle interactions. Physics of Fluids 18, 017101.

Liu, M.B., Liu, G.R., Zhou, L.W., Chang, J.Z., 2014. Dissipative Particle Dynamics (DPD): An Overview and Recent Developments. Archives of Computational Methods in Engineering 22, 529–556.

Majid, T., Griffin, D., Criss, Z., Jarpe, M., Pautler, R.G., 2015. Pharmocologic treatment with histone deacetylase 6 inhibitor (ACY-738) recovers Alzheimers disease phenotype in amyloid precursor protein/presenilin 1 (APP/PS1) mice. Alzheimers & Dementia: Translational Research & Clinical Interventions 1, 170–181.

Margulies, S., Thibault, L., Gennarelli, T., 1990. Physical model simulations of brain injury in the primate. Journal of Biomechanics Vol.23, 823–836.

Meaney, D.F, Ross, D.T., Winkelstein, B.A., Brasko, J., Goldstein, D., Bilston, L.B., Thibault, L.E., Gennarelli., T.A., 1994. Modification of the Cortical Impact Model To Produce Axonal Injury in the Rat Cerebral Cortex, Journal of Neurotruma 11 5.

Mehrafrooz, B., Shamloo, A., 2018. Mechanical differences between ATP and ADP actin states: A molecular dynamics study. Journal of Theoretical Biology 448, 94–103.

Misra, C., Ziff, E.B., 2005. EphB2 gets a GRIP on the dendritic arbor. Nature Neuroscience 8, 848–850.

Perez, F., Diamantopoulos, G.S., Stalder, R., Kreis, T.E., 1999. CLIP-170 Highlights Growing Microtubule Ends In Vivo. Cell 96, 517–527.

Peter, S., Mofrad M.R.K., 2012. Computational Modeling of Axonal Microtubule Bundles under Tension. Biophysical Journal 102 749–757.

Sanjith, S., 2011. Traumatic axonal injury in mild to moderate head injury – an illustrated review, Indian Journal of Neurotrauma (IJNT) 8, 71–76.

Shahinnejad, A., Haghpanahi, M., Farmanzad, F., 2013. Finite element analysis of axonal microtubule bundle under tension and torsion, Procedia Engineering 59, 16–24.

Shamloo, A., Manuchehrfar, F., Rafii-Tabar, H., 2015. A viscoelastic model for axonal microtubule rupture. Journal of Biomechanics 48, 1241–1247.

Shamloo, A., Mohammadaliha, N., Heilshorn, S.C., Bauer, A.L., 2015. A Comparative Study of Collagen Matrix Density Effect on Endothelial Sprout Formation Using Experimental and Computational Approaches. Annals of Biomedical Engineering 44, 929–941. doi:10.1007/s10439-015-1416-2

Shamloo, A., Mohammadaliha, N., Mohseni, M., 2015. Integrative Utilization of Microenvironments, Biomaterials and Computational Techniques for Advanced Tissue Engineering. Journal of Biotechnology 212, 71–89.

Shamloo, A., Mehrafrooz, B., 2018. Nanomechanics of actin filament: A molecular dynamics simulation. Cytoskeleton 75, 118–130.

Smith, A., Douglas, H., Meaney, DF., Shull, WH., 2003. Diffuse Axonal Injury in Head Trauma, Journal of Head Trauma Rehabilitation 18 307–316.

Smith, M.A., Blankman, E., Gardel, M. L., Luettjohann, L., Waterman, C. M., Beckerle, M. C., 2010. A Zyxin-Mediated Mechanism for Actin Stress Fiber Maintenance and Repair. Journal of Developmental Cell 19, 365–376.

Tang-Schomer, M.D., Patel, A.R., Baas, P.W., Smith, D.H., 2010. Mechanical breaking of microtubules in axons during dynamic stretch injury underlies delayed elasticity, microtubule disassembly, and axon degeneration. The FASEB Journal 24, 1401–1410.

Teixeira, C. A., Miranda, C. O., Sousa, V. F., Santos, T. E., Malheiroa, A. R., Solomonc, M., Gustavo, H., Maegawa, Brites P., Mendes Sousa, M, 2014. Early axonal loss accompanied by impaired endocytosis, abnormal axonal transport, and decreased microtubule stability occur in the model of Krabbe’s disease, Neurobiology of Disease 66, 92–103.

Williams, D.H., Levin, MA., Harvey, S., Eisenberg, Howard M. M.D., 1990. Mild Head Injury Classification, journal of Neurosurgery 27, 422–428.

Wu, J., Yuan, H., Li, L., Fan, K., Qian, S., Li, B., 2018. Viscoelastic shear lag model to predict the micromechanical behavior of tendon under dynamic tensile loading. Journal of Theoretical Biology 437, 202–213.

Wu, J.Y., Yuan, H., Li, L.Y., 2018. Mathematical modelling of axonal microtubule bundles under dynamic torsion. Applied Mathematics and Mechanics 39, 829–844.

Walstad, A., 2013. On deriving the Maxwellian velocity distribution. American Journal of Physics 81, 555–557.

Yu, W. Q., Baas, P. W., 1994. Changes in microtubule number and length during axon differentiation. Journal of Neuroscience 14, 2818–2829.

